# Population Consequences of Single-Cell Damage Dynamics: An Overlooked Role of Mortality Under Stress

**DOI:** 10.1101/2025.09.25.678661

**Authors:** Murat Tuğrul, Audrey M. Proenca, Ulrich K. Steiner

## Abstract

Microbial population growth arises from the survival and division of individual cells. However, under environmental stress, how reduced population fitness emerges from the single-cell dynamics remains poorly understood. Cellular aging and damage accumulation are often overlooked in linking these levels. Here, we use both theory and experiment to investigate how stochastic and asymmetric damage dynamics shape population outcomes. Theoretically, we apply a jump-diffusion damage model within a structured population framework to explore the roles of damage rate, noise, and partitioning asymmetry. Our theoretical analyses show that these parameters influence population growth both at equilibrium and during transient dynamics, suggesting that their cellular regulation may be a key strategy for sustaining population fitness. Experimentally, we expose Escherichia coli to glucose limitation and monitor stress responses using a RpoS fluorescent reporter, tracking both single-cell behavior in a microfluidic device combined with time-lapse fluorescence microscopy and population growth in a plate reader. Glucose limitation leads to the effects of elevated stress, reduced division, and increased mortality, which are consistently observed across scales. Using parameter estimates from single-cell data, our model deepens insights on population-level dynamics and highlights damage-noise driven, damage-dependent mortality as a key factor under stress. Together, these findings establish a quantitative framework linking intracellular stress to population fitness under environmental stress.

## INTRODUCTION

Microbial population growth arises from the survival and reproduction of individual cells. However, especially under environmental stress, how population-level outcomes emerge from single-cell level remains poorly understood. Cellular aging and damage accumulation are fundamentally intertwined with survival and reproduction (Kirkwood 1977), yet they are often overlooked in this individual-to-population scaling challenge. That is partly due to heterogeneous life trajectories: some cells may accumulate damage rapidly and die, some lucky ones escape high damage levels, while some others might partition damage asymmetrically during division and thereby rejuvenate themselves (Ackermann et al. 2007). This inherently stochastic and asymmetric nature of cellular damage makes it difficult to predict population-level outcomes. Still, understanding how single-cell aging and damage shape population dynamics is increasingly important, especially given the global relevance of microbial populations (Murray et al. 2022).

The recognition of aging in bacteria is relatively recent (Ackermann et al. 2003; Stewart et al. 2005). Unlike in multicellular eukaryotes, where aging is tied to mechanisms such as telomere shortening or stem cell exhaustion (López-Otín et al. 2023), bacterial aging is often linked to physical damage, such as misfolded proteins or protein aggregates, which accumulate stochastically over time and can be inherited asymmetrically during cell division (Lindner et al. 2008; Mortier et al. 2023; Proenca et al. 2024). This stochasticity and asymmetry can create heterogeneity in damage accumulation across lineages and may impact population fitness. However, establishing a direct link between molecular damage and aging remains challenging, as damage is diverse and its propagation across bacterial lineages is difficult to measure empirically (Steiner 2021). Given this complexity, abstract and overall quantifications are more useful and necessary, allowing integration of diverse damage types into a single aging framework (Gladyshev 2016).

On the empirical side, many studies use global stress-response regulators, such as the sigma factor RpoS in *E. coli* (Lange and Hengge-Aronis 1991), as proxy to intracellular stress or damage level. RpoS levels are informative even during exponential growth (Dong and Schellhorn 2009), and exhibit strong stochastic variation across single-cells (Patange et al. 2018). Given its central role, it is not surprising that studies have examined the influence of RpoS on longevity and senescence at the single-cell level. For instance, *rpoS* lacking mutant cells exhibit increased rates of aging (Yang et al. 2019). However, despite its widespread use in single-cell studies, the population-level consequences of RpoS activity remain poorly understood (Saint-Ruf et al. 2004; Bakshi et al. 2021), in part due to broader challenge of linking single-cells damage dynamics to populations. This gap limits our ability to understand and predict microbial population responses under stress conditions.

On the theoretical side, discussions on scaling aging to the population level are rooted in evolutionary theory (Partridge and Barton 1993; Charlesworth 1994), but applications to microbial systems remain limited. Existing models of bacterial aging that address population-level consequences often focus on the optimality of asymmetric damage partitioning to maximize fitness at equilibrium (Watve et al. 2006; Evans and Steinsaltz 2007; Blitvić and Fernandez 2020). However, these models typically neglect explicit dynamic processes and lack empirical confrontation. Other theoretical work has shown that stochastic variation in division rates can enhance population growth (Stukalin et al. 2013; Hashimoto et al. 2016), but these studies primarily focus on reproduction, often overlooking cell survival’s role in shaping population dynamics. In parallel, much of microbial population modeling has centered on resource limitation (Monod 1949) or density-dependent regulation (Contois 1959), primarily through their influence on division rates. While these are important, aging-related mortality is often ignored, which can play a crucial role under stress conditions. A more comprehensive framework is needed to link stochastic damage accumulation and asymmetric inheritance at the single-cell level to population-level outcomes, without ignoring mortality in populations.

In this study, we address the gap in understanding how single-cell aging and damage dynamics scale to shape population-level outcomes. We first lay out a theoretical framework for linking cellular damage to population dynamics. We then apply a mathematical model that incorporates two key features of bacterial aging: stochastic damage accumulation and asymmetric partitioning during cell division and examine their impact on population growth. Finally, we test the model using *Escherichia coli* exposed to glucose limitation, integrating single-cell and population-level measurements. Together, these steps offer a unified approach to understanding how damage processes at the single-cell level drive microbial population responses under stress.

## RESULTS

### Theoretical framework for scaling damage dynamics from single cells to populations

We begin by developing a general theoretical framework to capture how single-cell life trajectories comprising division and mortality events scale up to shape population dynamics. Under constant division and mortality rates, population growth follows a classical exponential model, i.e., *n*(*t*) = *n*_0_ *e*^λ*t*^, where *n*_0_ is the initial population size and λ = *r ln*2 − *h*_*o*_ represents the population growth rate, or the Malthusian fitness, determined by the division rate *r* and the mortality rate *h*_*o*_. This simple relationship gets more complex if aging is not negligible, and influences reproduction and mortality rates.

In our theoretical framework, aging is quantified by cellular damage, a single trait that conceptually represents the inverse of cellular vitality. We consider an extension of the McKendrick partial differential equation (M’Kendrick 1925; Keyfitz and Keyfitz 1997; Stukalin et al. 2013) to incorporate this single trait variable (Oizumi and Takada 2013) allowing us to describe how cellular damage *x* change over time *t* and age *a*, i.e.,

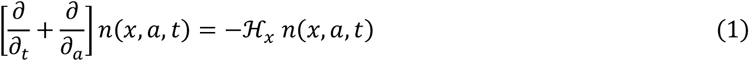

where *n*(*x, a, t*) is the population density of individuals with damage level *x* and age *a* at time *t*. The operator ℋ_*x*_ describes the statistical characteristics of stochastic damage trajectories including mortality events. Reproduction (newborns, i.e., *a* = 0) is captured via the boundary condition:

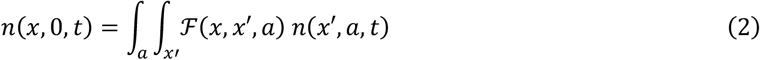

where ℱ(*x, x*′, *a*) is fecundity and damage inheritance from a parent with damage *x*′ and age *a*. The initial condition at t=0 defines the population by

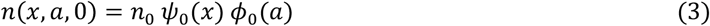

where *n*_0_, *ψ*_0_(*x*) and *ϕ*_0_(a) are initial population size, damage and age distributions, respectively. The population size at time t is calculated as

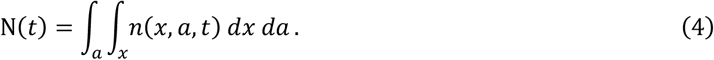

and the age and damage distributions at time t can be computed as *ϕ*(a, *t*) = ∫_*x*_ n(x, a, t)dx / N(t) and *ψ* (x, *t*) = ∫_*a*_ *n*(*x, a, t*)da / *N*(*t*), respectively. A long-term equilibrium is typically assumed by separating variables and expressing the solution in exponential form with a growth rate λ, i.e.,

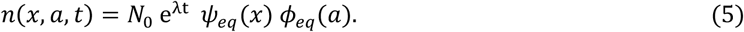

An analytical solution is usually difficult to obtain due to the complexities of ℋ_*x*_ and ℱ(*x, x*′, *a*), thereby, numerical solutions are favored.

Here, we use an age-stage-structured population, considering a projection matrix method as a known approximation to the McKendrick-type formulation (Keyfitz and Keyfitz 1997). Time is discretized as *t* = 0, Δ*t*, 2Δ*t*,.., age as *a* = 0, Δ*t*, 2Δ*t*, . . , *a*_*m*ax_ and trait as *x* = *x*_*min*._, *x*_*min*._ + Δ*x, x*_*min*._ + 2Δ*x*, …, *x*_*max*_. We define a population vector 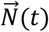 where each entry *N*_(*a,x*)_(*t*) corresponds to the count of individuals of age *a* and damage x at time t. The discrete dynamics over a single time step is then governed by

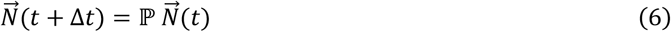

where each entry of the transition matrix ℙ_(*a*′,*x*′),(*a,x*)_ denotes the expected multiplicity of individuals projected from state (*a, x*) to state (*a*′, *x*′). For *a*′ > 0, ℙ_(*a*′,*x*′),(*a,x*)_ = *S*(*a, x*, Δ*t*) *K*(*x*′ | *x*) where K is the discretized transition kernel. For *a*′ = 0 (new born), ℙ_(*a*′,*x*′),(*a,x*)_ = *S*(*a, x*, Δ*t*) *b*(*a, x*) *K*_*inherit*_(*x*′ | *x*) where K is the discretized inheritance kernel and *b(a,x)* is the fecundity of the individual with state (a,x). The leading eigenvalue of ℙ corresponds to the population growth at equilibrium, i.e., λ = log(*m*ax(Eig(ℙ))). The corresponding eigenvector, when normalised, provides us the age and damage equilibrium distribution as an approximation to *ψ*_*eq*_(*x*) and *ϕ*_*eq*_(*a*). The dynamic equation in **Eq. (6)** with initial condition describes the population dynamics.

### Population growth under a jump-diffusion damage model: Roles of damage rate, noise and asymmetry

To investigate how single-cell damage dynamics give rise to population-level outcomes within the theoretical framework described above, we next consider a mathematical damage model capturing two key biological features: stochasticity (noise) in damage production and asymmetric damage partitioning during cell division. The minimal formulation within a stochastic differential equation framework is a jump-diffusion equation with constant parameters:

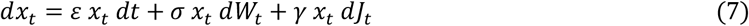

where ε is the deterministic damage production rate, and σ represents noise strength (modeled with a Wiener process *W*_*o*_) and γ is the asymmetry in damage partitioning during division events (modeled with a jump process defined by *J*_*o*_). Here γ = 0 indicates symmetric division, and 0 < γ < 1 reflects mother (old pole) cell lineages receiving more damage than the daughter (new pole) cell. While this minimal model abstracts biological complexity, it incorporates the key features of stochasticity and asymmetry, and offers analytical tractability which is an important property for theoretical and data analyses.

In our previous work (Tuğrul and Steiner 2025), we examined how this particular damage dynamics shapes lifespan statistics in a cohort of single-cell mother lineages with fixed initial damage *x*_*c*_ and a critical damage level *x*_*c*_ defining the cell death. We obtained an approximate solution by assuming a Gaussian distributed division rate with a mean rate r and standard deviation *σ*_*r*_. This approximate solution facilitates analytical expressions for the transition probability density along mother cell lineage from an initial damage *x*_*c*_ to *x* within a time interval t as

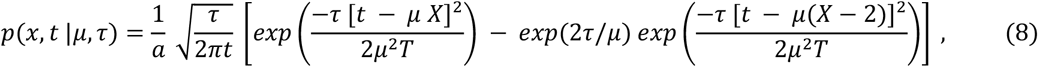

and the lifespan distribution of individual mother cell lineages as

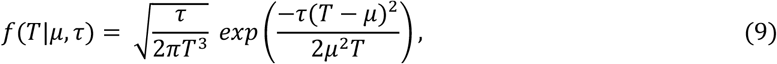

which is an inverse Gaussian distribution where the mean parameter

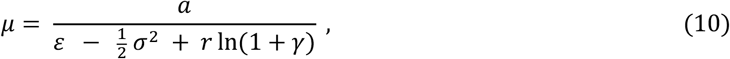

and the shape parameter

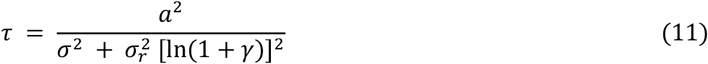

serve as projection from cellular damage parameters to demographic parameters, where *a* = ln *x*_*c*_/*x*_*o*_ serves as a time scaling factor and 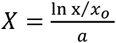 is used for a compactness. This approximation corresponds to a Geometric Brownian Motion in the form of 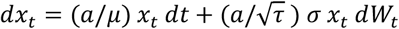. The survival probability can also be expressed analytically using 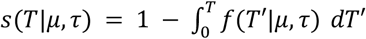 and we obtain

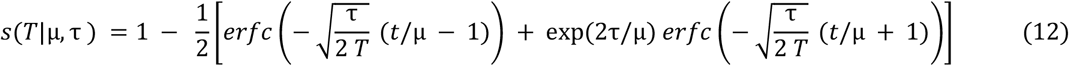

An external (i.e., damage-independent) killing, in terms of a Poisson process of a rate *h*_*o*_, can easily be integrated into this model by modifying the survival probability in **Eq. (12)** as

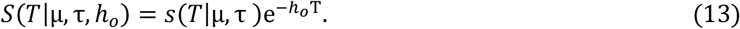

These expressions form a tractable mapping from damage dynamics at single-cell level to demographic parameters such as lifespan distributions (**Eq. (9)**) and survival probability (**Eqs. (12-13)**). Next, we use their discretized versions to construct the transition matrix in **Eq. (6)**. To finalize the matrix construction’s inheritance kernel, we assume that all 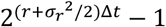 offspring (daughter cell lineages) are born at the middle time point Δ*t*/2.

Figure 1. illustrates the population-level consequences of the theoretical jump-diffusion damage model, using both exact simulations of **Eq. (7)** in a growing population and the above explained approximation for population dynamics (**Eqs. 6, 8-13**). **Fig. 1B–E** compare two scenarios of damage dynamics, low-stress and high-stress, which we achieved by increasing both the damage production rate and noise intensity, while keeping other parameters, including the division rate, constant. To isolate the effects of damage, external killing was also set to zero in this theoretical analysis. Under high-stress conditions, cells accumulate damage more rapidly and variably, resulting in population-wide higher damage levels, as seen in the example simulation of the damage dynamics throughout the cell population in **Fig. 1B**.

**Figure 1.**
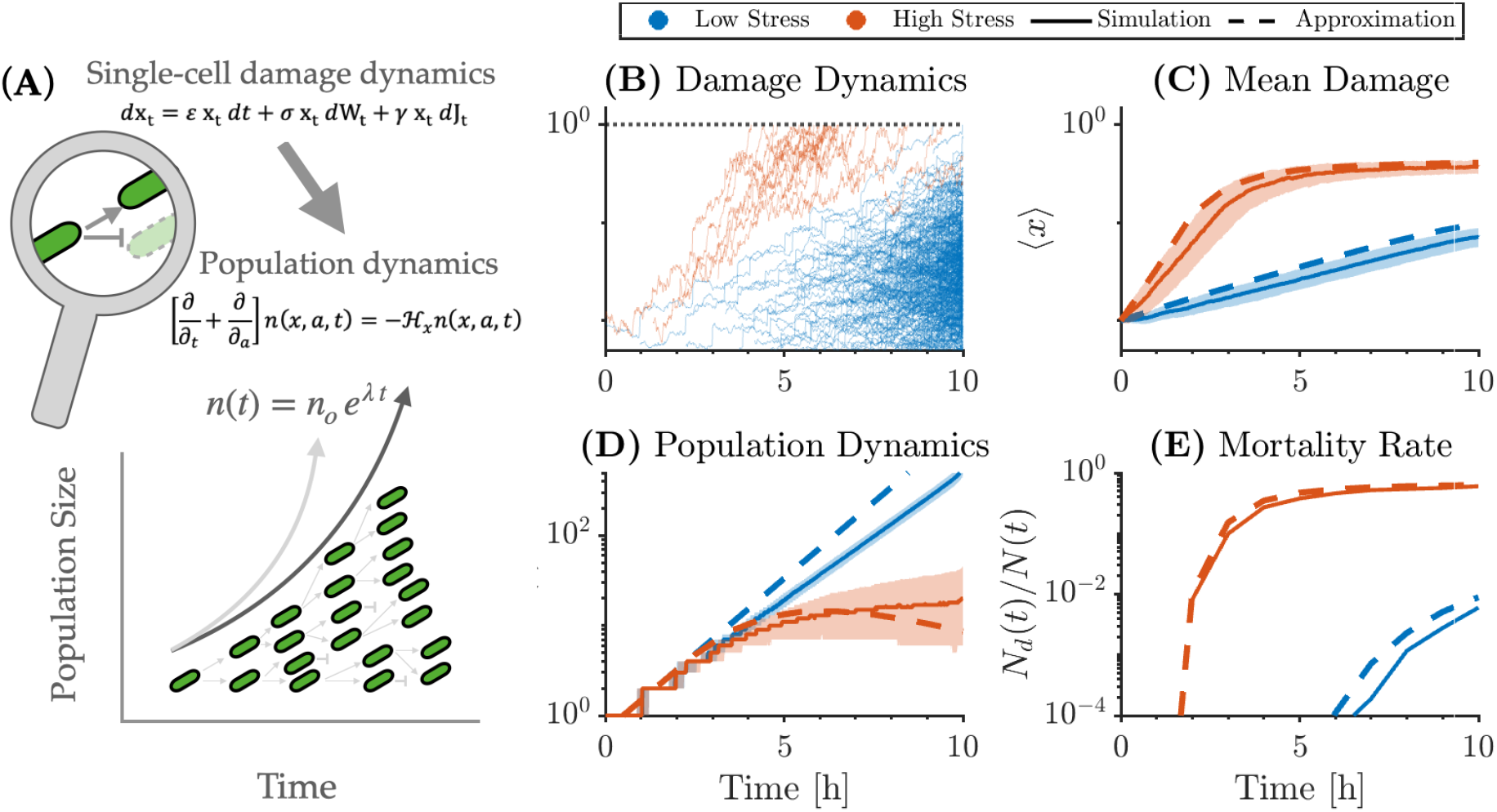
Population-level consequences of the jump-diffusion cellular damage model. (A) Schematic of the modeling framework linking single-cell damage dynamics to population dynamics. (B–E) Comparison of low-(blue) and high-stress (red) conditions simulated by increasing damage production and noise (i.e., *ε* = 0.1 vs *ε* = 1.0 while keeping *σ*^2^/*ε* = 0.5, *γ* = 0.5, *r* = 1.0, *σ*_*r*_ = 0.25, *x*_*o*_ = 0.01, *x*_*c*_ = 1.0) (B) Example damage trajectories along each cell lineages in a growing population. Our age-damage structured population approximation (dashed curves) closely reproduces the results of the exact simulations (solid curves) of damage dynamics for these parameter selections, as seen in (C) mean damage in population across time, (D) population size dynamics, and (E) mortality rate over time.

Notably, the average damage level approaches the critical threshold defining the cell death more quickly under stress (**Fig. 1C**). This leads to an earlier and more intense number of mortality events or rates (**Fig. 1E**). This increase in damage-induced mortalities significantly reduces population growth, as reflected by the slowing down in the population growth rate in **Fig. 1D**. This growth reduction arises solely from damage-linked mortality, as reproduction rates remain unaffected and no additional constraints, such as density dependence, are included in the model. Our approximate age-damage structured population model closely follows the exact simulation results, generally capturing the time scaling of population size, population-level mortality rates and mean damage levels.

**Figure 2A** expands the analysis by using the matrix method and shows how population growth responds to varying damage noise and damage partitioning asymmetry across a range of damage production rates representing an increasing level of external stress. Recall that we report equilibrium population growth rates by calculating the logarithm of the leading eigenvalue of the projection matrix. To facilitate interpretation across conditions and enable comparison with cell division dynamics, the resulting exponential growth rates are converted to doubling rates and scaled by the mean cell division rate. Under low damage production (low-stress environments), as expected, population growth is largely unaffected by noise or asymmetry. In this regime, damage-induced mortality is rare, and growth is determined almost entirely by the division events. As damage production increases, differences between parameter settings become pronounced. At increased but still moderate stress levels, intermediate asymmetry in damage partitioning leads to the greatest reduction in population growth. In contrast, extreme asymmetry values, which are either near-symmetric division (*γ* → 0) or highly asymmetric division favoring damage segregation into one cell (*γ* → 1) help maintain higher population growth rates by either diluting damage across the population or eliminating it from the population with “garbage collector” dying mother cells. At even higher damage production rates, the influence of noise becomes apparent. Increased stochasticity allows for rare “lucky” cell trajectories that avoid high damage, supporting the survival and persistence of the population under otherwise lethal conditions. In this regime, highly asymmetric damage partitioning (*γ* → 1) emerges as the most optimal strategy, effectively isolating damage into mother lineages, but *γ* → 0 solution disappears. Thus, both asymmetry and noise play context-dependent roles in shaping population-level fitness under stress.

**Figure 2.**
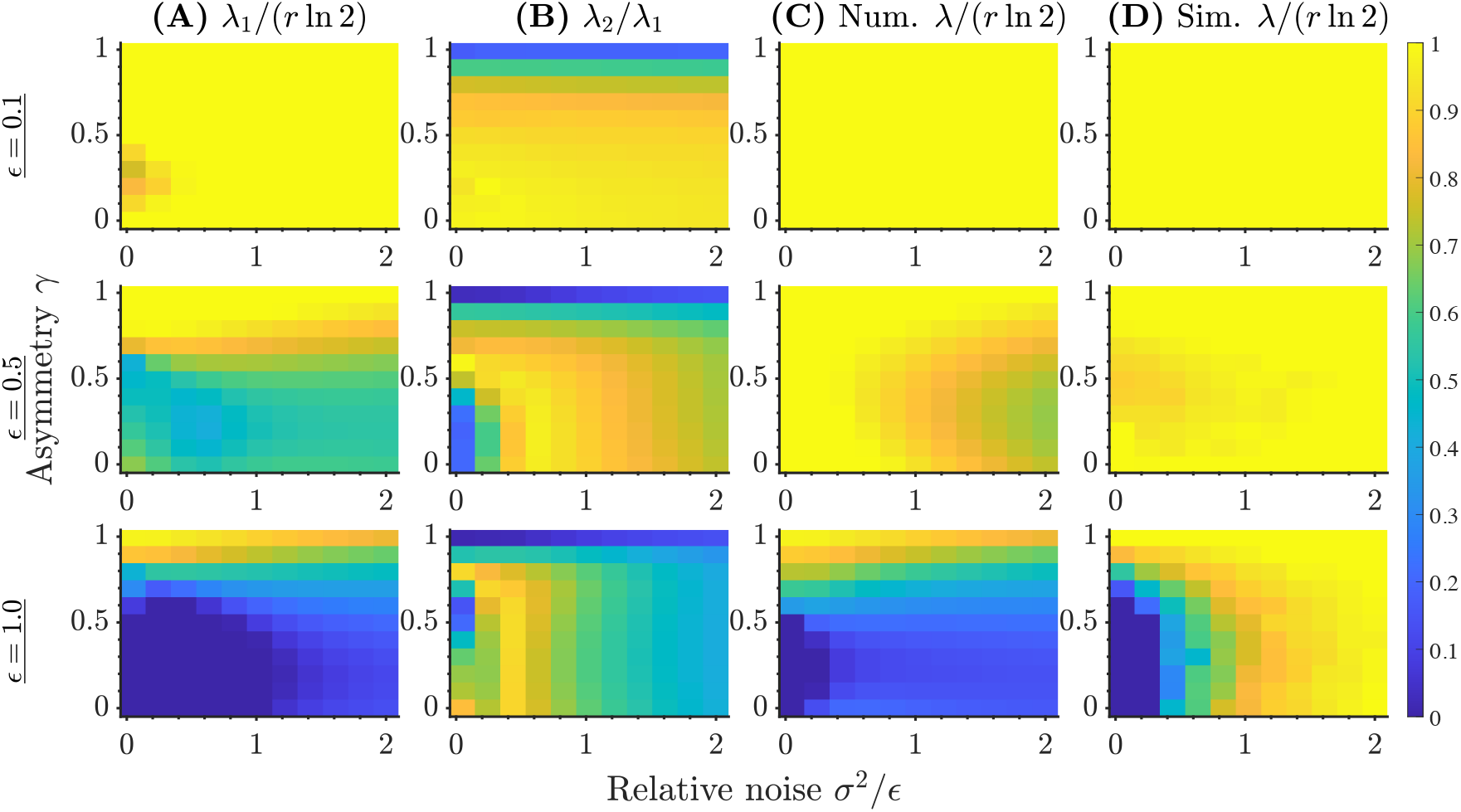
Population growth rate at equilibrium, convergence and dynamic behavior. **(A)** Heatmaps showing the equilibrium population growth (doubling) rate (scaled with division rate *r*) across varying damage noise and damage partitioning asymmetry for three increasing damage accumulation rates. (B) Ratio of the two leading eigenvalues of the projection matrix, reported as a proxy for the convergence rate toward the equilibrium growth solution. (C-D) Numerical replication of (A) to estimate the population growth rates over the 4–7 h window by (C) iterating the projection matrix and (D) performing exact simulations of damage dynamics starting from a low-damage initial state and fitting exponential growth. Results show the slow approach to equilibrium solutions at certain parameter settings.

The results above are based on the equilibrium assumption, reflecting long-term population dynamics. In principle, populations may approach this equilibrium slowly, depending on initial conditions. In **Figure 2B**, we report the ratio of the two leading eigenvalues of the projection matrix as an indicator of the convergence rate. We also replicate the analysis in **Fig. 2C** by numerically iterating the projection matrix and in **Fig. 2D** by simulating the exact damage dynamics in a growing population, starting from a low-damage (“healthy”) initial state and fitting an exponential curve to the resulting population size over the 4–7-hour time window. These analyses confirm that convergence can be slow under certain parameter settings, and thus equilibrium-based results, which is widely assumed in the literature, should be interpreted with caution. When possible, explicit dynamical modeling should be used to complement asymptotic analyses. Whether natural microbial populations have evolved mechanisms to tune damage characteristics for optimal population performance remains an open question and is beyond the scope of this study. To explore empirical correspondence with the theoretical model developed thus far, we present a simple experimental design and analysis in the next section.

### Experimental Application: Glucose-limitation stress in E. coli at single-cell and population levels

To investigate how external stress conditions influence intrinsic cellular stress and fitness across scales of single-cell and population, we performed a simple experiment using E. coli cells carrying a fluorescent transcriptional reporter for RpoS promoter activity (PrpoS-GFP) (Zaslaver et al. 2006). This reporter serves as a proxy for overall cellular stress response, and in our previous work, we showed that RpoS activity is inversely correlated with cell elongation rate, an important component of cellular fitness (Proenca et al. 2024). In this study, we examine how E. coli responds to glucose-limitation stress (0.0% glucose) in comparison to standard level (0.4% glucose) under minimal medium M9 in both single-cell and population levels.

To collect single-cell data, we used a microfluidic device combined with time-lapse fluorescence microscopy, known as the “mother machine” (MM) (Wang et al. 2010). It maintains a continuous flow of fresh medium through main channels, feeding narrow side channels that trap mother cells inheriting the older pole during division. This setup mimics conditions of exponential population growth by providing constant nutrients and removing waste and allowing observation of single-cell dynamics. Using a machine learning-based pipeline (Lugagne et al. 2020), single-cells were segmented and tracked, yielding *979* lineages under standard glucose conditions and *332* under glucose-limitation stress (right-censored: *647* and *202*, respectively). This allowed us to quantify physiological traits (e.g., cell size, division timing), RpoS promoter activity via GFP fluorescence, and membrane permeability using propidium iodide (PI) as a red-fluorescent marker of cell lysis up to *72* hours with *4* min intervals.

We first examined two key components of individual fitness in single *E. coli* cells: division and mortality rates. Division rates were quantified using a statistical approach based on the distribution of division counts within 10-hour windows. The distributions were bimodal, separating slow or dormant cells from actively dividing ones. In both conditions, reproductively active cells maintained stable division rates over time (**Fig. 3B**), consistent with previous findings (Wang et al. 2010), with a slight increase over time likely reflecting adaptation. Under glucose limitation, the distribution shifted modestly lower, indicating slower growth in comparison to glucose-rich environment (mean division rate: *1*.*05* vs *1*.*20* div/h in early times; and *1*.*20* vs *1*.*60* div/h in later times).

**Figure 3.**
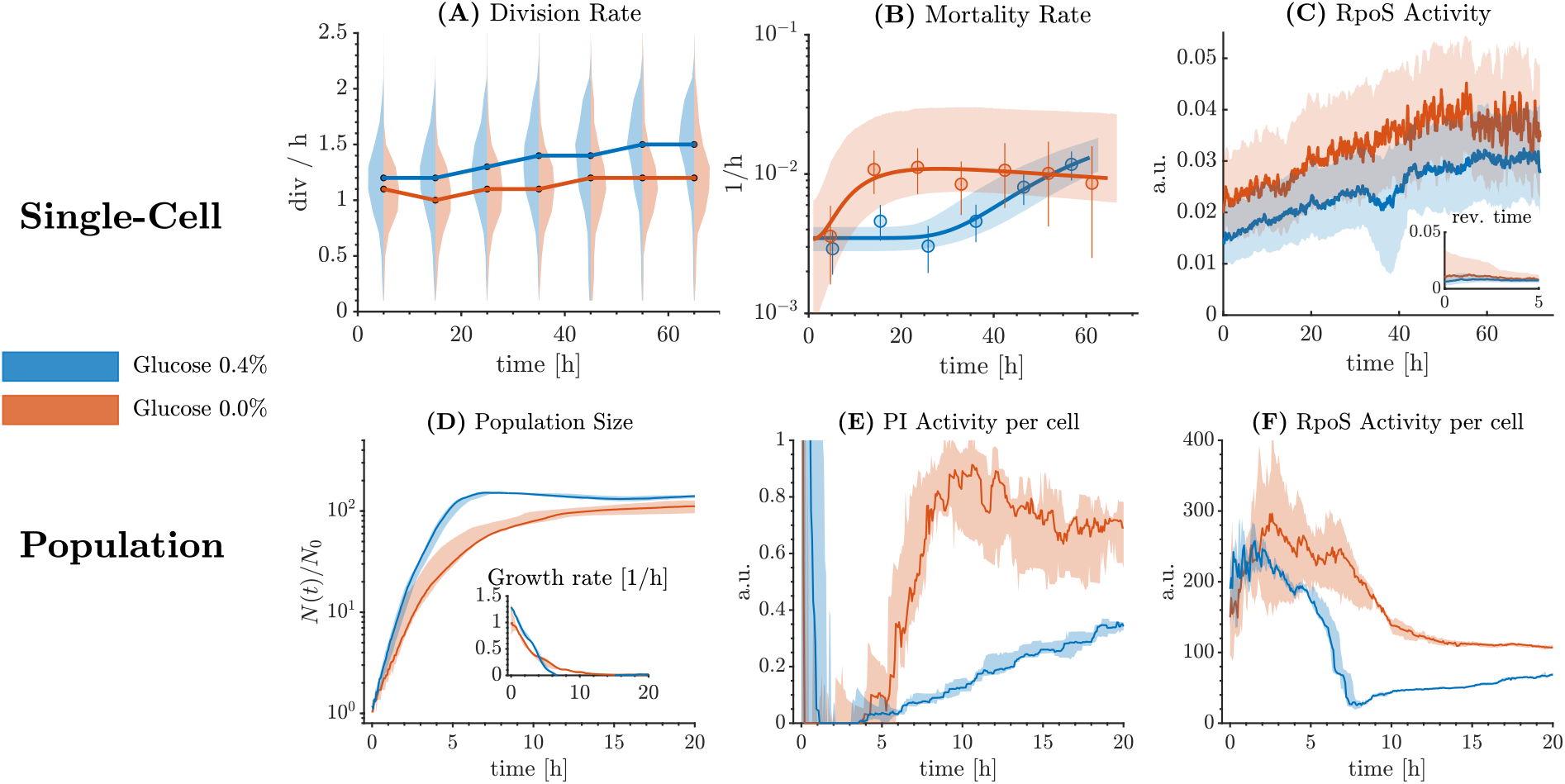
Experimental analysis of single-cell and population responses to glucose limitation stress in E. coli. Single-cell microfluidics (mother machine) and population-level plate reader measurements under glucose-rich (blue) and glucose-limited (red) M9 minimal media. (A) Division rate dynamics estimated from active single-cell lineages (*r > 0*.*25)* are presented as half violin plots and the curves follow the medians. (B) Mortality rate statistics estimated from growth arrest and cell lysis (PI) signals. The circles with error bars show the experimental data whereas solid curve and shading are for the damage model fitting. (C) RpoS activity over time in single cells; inset shows reverse-time alignment from death/arrest. (D) Plate reader OD measurement is converted into population size dynamics normalized by initial size, i.e., N(t)/N_0_. Inset shows exponential growth rate over time. (E) Per-cell PI fluorescence as a proxy for lysis and mortality. (F) Per-cell RpoS activity reflecting stress response.

To estimate fitness more fully, we also quantified single-cell mortality rates. Cell death was detected using PI fluorescence (lysis marker), as well as using a stringent growth arrest criterion, defined as no exponential elongation rate exceeding of an exponential rate 0.05 px/h during any 4-minute interval over a one-hour period. Unlike division rates, mortality showed a pronounced divergence between conditions (**Fig. 3B**). Both conditions consistently began with a low basal mortality rate of ∼0.003/h. This is presumably corresponding to the damage-independent killing rate due the shared initial environment in our experimental set-up, where cells originate from an exponentially growing population culture and are unlikely to have accumulated any damage. However, in glucose-limited cells, mortality rose sharply within a few hours and plateaued around ∼0.01/h, whereas in glucose-rich conditions, mortality remained low for ∼*24* hours before gradually increasing. This divergence is consistent with our prior analysis of various single-cell datasets comparing wild-type strains to interventions (e.g. mutants) that are possibly stress-inducing (Tuğrul and Steiner 2025). Overall, these findings suggest that mortality plays a critical role alongside reproduction in shaping the fitness of single bacterial cells under stress.

To understand whether mortality is linked to internal stress, we analyzed RpoS reporter dynamics. Total RpoS signal was measured within the cell area and known GFP decay rate was used to estimate RpoS activity (see Methods). As shown in **Fig. 3C**, cells under glucose limitation exhibited higher RpoS activity over time, with both conditions showing similar rates of increase, but interestingly the glucose-limited case caused a higher cell-to-cell variability. Supporting along this, reverse-time alignment of single-cell trajectories (**Fig. 3C, inset**) revealed even more pronounced variation in upregulation of RpoS shortly before cell lysis and growth arrest, suggesting a strong association between stochasticity in stress response activation, division arrest, and mortality.

Next, we addressed whether *E. coli* populations in bulk culture reflect stress-induced changes in growth and mortality. We used the identical growth medium as in the single-cell experiments and performed two biological replicates, each with three technical replicates per condition in a 96-well plate reader. We measured OD (optical density at 600nm), GFP (RpoS activity), and PI (cell death) fluorescence over time. OD values were background-subtracted and aligned by shifting curves to a common threshold (OD = 0.004), which also served to normalize the OD measurements to estimate relative population size as N(t)/N_0_ (**Fig. 3D**). Note that we also used the cell size differences observed in single-cell level in this population size calculation. Population growth began with the exponential rates consistent with the above reported division rates in single cells but declined over time gradually (**Fig. 3D, inset**). In glucose-rich media, populations reached carrying capacity by ∼*6* hours, while under glucose limitation, growth was slower, and saturation occurred only after ∼10 hours. Importantly, *E. coli* cells show a significant reduction in population growth under glucose limitation, in agreement with single cells. Notably, in the absence of glucose, *E. coli* cells depend on the casamino acids in the media for growth. This was confirmed by our population-level control experiments (not shown), resulting in no growth without casamino acids, which is consistent with single-cell observations reported by others (Gervais et al. 2024).

Can the observed reduction in population growth be explained solely by nutrient or metabolic shifts affecting only division rates, or do stress-induced mortalities also contribute? To address this, we analyzed PI and RpoS signals from plate reader assays. PI, a proxy for cell lysis and death, was normalized by estimated population size. The obtained PI per cell showed a clear rise around 4th hour in both conditions where the glucose limitation case showed a rapid increase in short time, indicating elevated mortality under stress (**Fig. 3E**). Similarly, per-cell RpoS activity (normalized similarly) increased in both conditions but was higher under no-glucose stress (**Fig. 3F**). They peaked around in the 3–5-hour band and start decreasing afterwards, likely due to death and elimination of highly stressed cells. These population-level patterns in growth, stress, and death aligned with our single-cell observations (**Fig. 3A-C**).

### Connecting single-cell and population-level dynamics through a damage model

The alignment between single-cell measurements and population-level outcomes suggests the presence of a shared aging or damage-driven process operating across scales. To explore this possibility, we employed our theoretical model of cellular damage where the model’s key parameters were inferred directly from single-cell data to investigate the consistency in bridging to population dynamics.

Division rates were modeled as Gaussian-distributed with means of 1.20 vs. 1.60 divisions per hour and standard deviations of 0.25 vs. 0.33 div/h under glucose-limited and glucose-rich conditions, respectively, based on measurements at late (adapted) time in single-cell experiments (**Fig. 3A**). Mortality parameters were estimated using lifetime data from the same experiments, incorporating right censoring information for cells that were alive at the end of the experiment, and a 3-hour left truncation to account for the time between cell loading and imaging onset in our experiments, which allows for cells to adjusts to the microfluidic environment. Maximum likelihood estimation yielded the following parameter estimates (95% confidence intervals in parentheses):

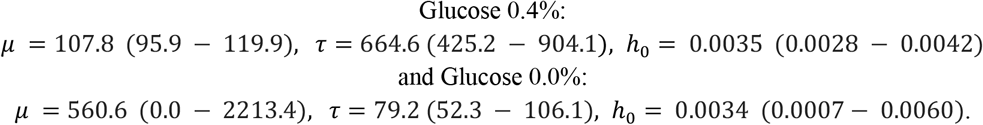

The fitted mortality rate curves and confidence bands are shown in **Fig. 3B**. Notably, both conditions yield consistently similar estimates for the basal mortality rate *h*_0_, likely reflecting shared environmental conditions in the mother machine during the early, low-damage phase. In contrast, the index of dispersion, i.e., *D* = *μ*^2^/ *τ*, a damage-free measure of lifetime variability and a key indicator of underlying damage process in our model (Tuğrul and Steiner 2025), differed by two orders of magnitude: ∼*1/36* under glucose-rich and ∼*36* under glucose-limited conditions. These values suggest fundamentally different damage regimes or mechanisms, indicating underdispersed and overdispersed lifetimes relative to a Poisson process.

Moreover, we used the RpoS information from the single-cell experiments and analyzed the partitioning asymmetry of total RpoS at division which interestingly remained stable over time, with a median value of *γ* = 0.06 (with a 25-75% CI of ∼0.02-0.12, **Fig. 4B**). This range is consistent with previous estimates based on size differences arising from protein aggregation (Wagener et al. 2024). Finally, we highlight that the RpoS activity trajectories (**Fig. 3C)** exhibit greater variability under glucose limitation stress but increase at similar rate across conditions. This observation supports the interpretation of a higher noise parameter *σ* under stress, with a similar damage rate *ε*. Note that this is consistent with the inferred values of *μ* and *τ* (**Eqs. (10–11**)), suggesting the importance of stochasticity in shaping the damage trajectories.

**Figure 4.**
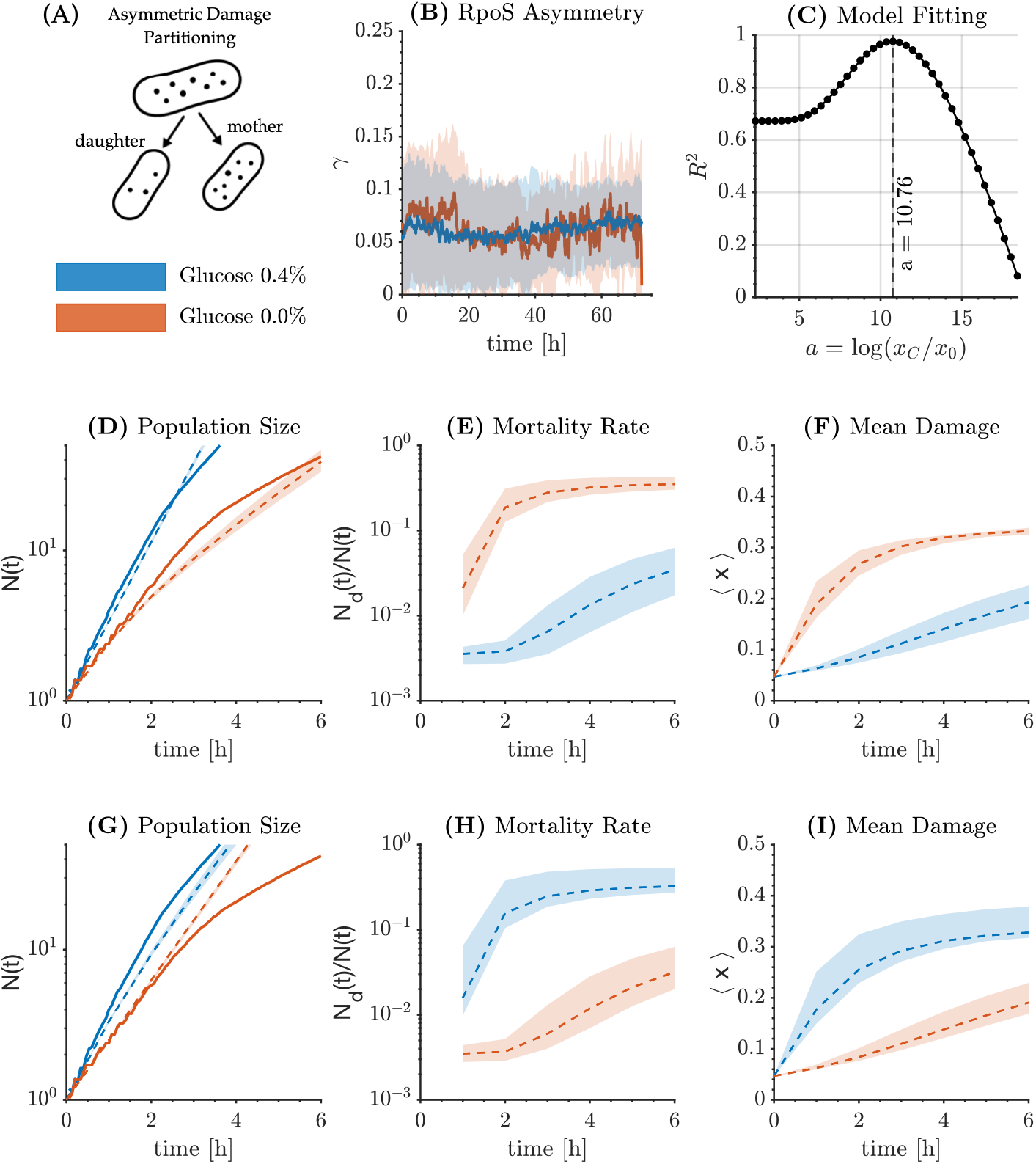
Fitting the damage-based population model to experimental data using single-cell parameter estimates. (A) Schematic illustrating asymmetric damage partitioning at cell division. (B-H) Glucose-rich (blue) and glucose-limited (red). (B) Distribution of RpoS asymmetry during divisions in single-cell data; the asymmetry parameter γ remains stable over time (median γ = 0.06; 25–75% CI ≈ 0.02– 0.12). (C) Model fit (R^2^) across the scaling parameter *a* = *log*(*xC* / *x*0) for the glucose-limited data, showing a clear optimum at a = 10.76. (D-I) Solid curves represent the median of experimental data. Dashed curves with shading represent the simulated model prediction’s median and CI (%75-25 for population size and %97.5-2.5 bands for mortality rate and mean damage). (D) Simulated population size dynamics using fitted parameters compared to experimental growth. (E) Predicted mortality rates from the damage-based population model show rapid increase under stress and gradual rise in the no-stress condition. (F) Mean cellular damage levels over time from the model, reproducing elevated damage under glucose limitation. (G-H) To test the contribution of inferred damage dynamics, we swapped all parameters between conditions except division-related ones and repeated the analyses in (D-F). (G) The resulting simulation failed to reproduce the observed population growth slowdown under glucose limitation. (H-I) The swap also reversed the predicted mortality rates (H) and mean damage levels (I) between conditions. Overall, damage-related parameters are essential to reproduce stress-induced population slowdown.

Using the parameter estimates derived from single-cell data under glucose-rich (non-stress) and glucose-limited (stress) environmental conditions, we projected damage dynamics at individual level to the population level by implementing an age-damage structured population model (**Eq. 6)**. In this model, the population is represented as a distribution of individuals across discrete damage states and ages. At each time step, individuals can divide or die based on their damage level, and damage is updated stochastically through accumulation and partitioning during division (**Eqs. 7-13)**. The only remaining free parameter was the dimensionless scaling constant *a* = ln *x*_*c*_/*x*_*o*_. We calibrated this parameter using the glucose-limited data, which showed a clear peak corresponding to the best-fitting value across conditions (at a = 10.76 with R^2^ = 0.98, **Fig. 4C**) and used this value for both conditions. Model fitting was restricted to the early phase of population growth to ensure the dynamics remained in the exponential regime prior to the onset of possible density-dependent regulation. A cut-off can be defined in the logistic growth framework, in which the inflection point is at half carrying capacity and corresponds to the time window up to half of the observed maximum population size. In our case, this is around *N*/*N*_0_∼50, and corresponds to the first *∼6* hours.

Next, we simulated population dynamics using a projection matrix built from single-cell parameter estimates and their confidence intervals. Starting from an initial damage state (*x*_*o*_ = Δx = 0.01, *x*_*C*_ = 1.00), we iterated population growth Δt = 1 hour intervals and compared it to experimental trajectories from the plate reader. The model closely captured the divergence between conditions and the slowed growth under glucose limitation (**Fig. 4D**), supporting that single-cell damage and aging shape population-level outcomes in stress environments. Using this framework, we also estimated average mortality rates as the ratio of death events to total individuals (**Fig. 4E**). While not calibrated to match PI-based measurements quantitatively, the theoretical trends of rapid mortality rise under stress and gradual increase in no-stress qualitatively aligned with experimental results. Finally, mean cellular damage levels calculated from the matrix method qualitatively captured elevated damage in glucose-limited populations at initial times (**Fig. 4F**), though they did not reflect the sharp decline observed experimentally at later times.

To further assess the role of damage-related parameters versus division rates, we swapped all parameters between conditions except those related to division. This resulted in the model failing to reproduce the full slowdown in population growth, confirming the contribution of damage accumulation and mortality, rather than slowed division alone (**Fig. 4H**). This swapping also reversed the predicted mortality rates and mean damage levels between conditions. In other words, incorporating damage-driven mortality, which was directly inferred from single-cell data, was essential to match the experimental trends at population-level. Considering division rate changes alone was insufficient. These findings highlight that mortality, driven by accumulated damage, is a key determinant of population dynamics under stress, beyond the effects of reduced reproduction.

## DISCUSSION

Understanding how single-cell aging and damage shape population-level outcomes remains a fundamental challenge, especially under environmental stress where stochasticity and asymmetry influence cell fates. To address this, we first presented a structured population framework and then applied it to apply a jump-diffusion model of damage accumulation, which was previously analyzed in a cohort context (Tuğrul and Steiner 2025). We thereby investigated how damage rate, stochastic noise, and partitioning asymmetry shape population growth. We revealed that under high damage rates, noise and asymmetry become increasingly critical. This can indicate that their regulation may serve as strategies to sustain population fitness. Unlike previous studies suggesting an optimal intermediate asymmetry level (Evans and Steinsaltz 2007; Watve et al. 2006; Blitvić and Fernandez 2020), our results, based on both simulations and analytical approximations, indicate that optimality is context-dependent and shaped by the interplay of damage rate and noise. These differences may stem from distinct modeling assumptions across studies. While our model offers analytical tractability and general insights, it represents one abstraction of damage dynamics. Future work may explore alternative formulations that incorporate larger biophysical detail while preserving the ability to scale across biological levels.

We quantitatively linked intracellular damage accumulation to reductions in microbial population fitness by integrating single-cell and population-level experiments of *E. coli* under glucose limitation with our theoretical framework. This combined approach demonstrates how stress-induced cellular aging shapes population dynamics. Importantly, our findings underscore the importance of mortality in shaping the population growth under stress conditions. Traditional approaches have largely emphasized modulation of division rates through density dependence, resource limitation. or metabolic switching. While important, these mechanisms alone cannot account for all the population dynamics observed under glucose limitation. Our results demonstrate that damage-induced mortality, which is inferred from single-cell data and incorporated into our model, is essential to capture the observed slowdown in population growth. Theoretical predictions using only division rate changes were insufficient. Aging and cellular damage dynamics need to be more deeply integrated into our general understanding of microbial populations, particularly under stress conditions.

The theoretical framework presented here provides a flexible platform for scaling intracellular processes to population consequences. Future extensions can include modeling different stochastic differential equation formulations for cellular damage, multi-trait damage dynamics, feedback between damage level and division rates. Additionally, integrating our theoretical framework into evolutionary models could help uncover how stress-induced damage, and the regulation of its stochastic and asymmetric features, shape the evolution of microbial populations, particularly under selective pressures like antimicrobial exposure (Mitosch et al. 2017). Our work also contributes to a broader understanding of how stress responses manifest at different biological scales. We show that intracellular stress dynamics measured with RpoS activity at the single-cell level can inform population-scale trends in both mortality and growth. Additional molecular indicators of damage or stress response will improve to deepen mechanistic insight (Gervais et al. 2024).

Together, our study highlights the need for integrating stochastic and asymmetric cellular aging and damage processes into microbial population models. By bridging single-cell dynamics with population-level consequences, we provide a foundation for more accurate predictions of microbial populations under environmental stress.

## ACKNOWLEDGEMENTS

We thank Shripad Tuljapurkar and Ryo Ouzumi for discussions. MT was funded by the Marie Skłodowska-Curie Actions’ European Postdoctoral Fellowship (grant 101069035), UKS was funded by the Heisenberg Programme of the German Research Foundation (grant 430170797), and AMP was funded by the Humboldt Research Fellowship (Alexander von Humboldt Foundation, Germany) and the Rising Star Research Fellowship (FU Berlin, Germany). The authors would like to thank the HPC Service of FUB-IT, Freie Universität Berlin, for computing time.

## MATERIALS AND METHODS

### Strains and Media

The E. coli K-12 strain MG1655 used in this study carried a plasmid-based GFP reporter to monitor RpoS promoter activity (PrpoS-GFP) (Zaslaver et al. 2006). Half-life of GFP is reported as 2.8h +/-0.7h, which corresponds to an exponential decay rate of 0.25 +/-0.06 per hour (Halter et al. 2007). Half-life of Propidium Iodide activity is not known, therefore based on our own observation, we set its decay rate as 1.0 per hour. Cultures were grown in M9 minimal medium containing 1xM9 salts, 2 mM MgSO_4_, and 0.1 mM CaCl_2_, supplemented with 0.2% Casamino Acids. Glucose was added at 0.4% for standard growth conditions and at 0.0% to induce carbon starvation stress. To prevent cell adhesion within microfluidic devices, 0.075% Tween-20 (Sigma-Aldrich) was included in the media, along with 1 µg/ml Propidium Iodide (PI) as a marker for cell lysis. To ensure comparability, the same media composition was used for both single-cell and batch culture measurements. All experiments were conducted at 37°C.

### Microfluidic device design and fabrication

Single-cell imaging was performed using a microfluidic device, named “mother machine” (Wang et al., 2010). It consists of four parallel flow channels (1 cm × 80 µm × 10 µm), each containing ∼1000 growth microchambers (25 µm × 0.8 µm × 1.2 µm) for capturing individual mother cell lineages. The master mold was fabricated via photolithography, and epoxy replicas were used for soft lithography casting with Sylgard 184 PDMS (Dow Corning, USA). PDMS casts were degassed, cured at 80°C for 1 h, and punched with 0.5 mm biopsy tools to form inlet and outlet ports. Devices were cleaned, bonded to 24 × 40 mm coverslips via plasma treatment, and cured overnight at 60°C. Prior to use, channels were plasma-activated and treated with 20% polyethylene glycol (PEG) for 1 h to minimize cell adhesion.

### Microfluidic device loading and setup

Overnight *E. coli* cultures were diluted into 15 ml M9 medium and grown for 2 hours at 37°C to reach exponential phase (OD_600_ = 0.4–0.6). Cell cultures were centrifuged at 4000 rpm for 10 min, resuspended in 200 µl M9 supplemented with 0.075% Tween-20 and 1 µg/ml PI. Then the cells were loaded into the microfluidic devices via the outlet ports and centrifuged at 1500 rpm for 8 min to fill the growth microchambers. After confirming proper loading, media flow was initiated at 200 µl/h using a peristaltic pump (Ole Dich, Denmark). Tygon tubing was replaced with flexible tubing (Masterflex Tygon E-3603, VWR) through the pump. Devices were maintained at 37°C in a temperature-controlled microscopy chamber (PeCon TempController 2000-1).

### Batch culture measurements using a 96-well plate reader

To complement single-cell analyses, population-level growth and fluorescence measurements were conducted in 96-well plates under the same media and temperature conditions. Overnight E. coli MG1655 cultures carrying PrpoS-GFP were diluted 1:100 into M9 minimal medium (with or without glucose) in a final volume of 200 µl per well. Cultures were prepared using the same protocol as for microfluidic experiments to ensure consistency across experimental platforms. Plates were closed with heads to minimize evaporation and loaded into a Biotek Synergy HT Multi-Mode Microplate reader maintained at 37°C. Optical density at 600 nm (OD_600_), GFP fluorescence (excitation: 485 nm; emission: 528 nm) and RFP fluorescence for propidium iodide (excitation: 540 nm; emission:590 nm) were recorded every 4 minutes for up to 24 hours. Plates were shaken before and after each measurement cycle to ensure culture homogeneity. Each condition was conducted in technical and biological triplicates, and background signals were subtracted using wells containing media only.

### Microscopy imaging

Time-lapse imaging was conducted using a Nikon Eclipse Ti2 inverted microscope with a motorized stage and Perfect Focus System (PFS), using 100x oil objective, and a DS-Qi2 camera. Image acquisition (phase contrast and fluorescence) was automated via NIS Elements software, with frames captured every 4 minutes.

### Image processing

Image preprocessing was performed in ImageJ for chromatic shift correction and background subtraction using a rolling ball algorithm (radius = 20 px, sliding paraboloid). Segmentation and tracking were conducted using the DeLTA (v1) machine learning pipeline (Lugagne et al. 2020) on the HPC cluster at FU Berlin. Phase contrast images were used for segmentation, and extracted features included cell length, area, division timing, and mean fluorescence intensity.

### Data analysis

All data analysis was performed using MATLAB (tested in versions R2021b and R2024b, MathWorks). Custom scripts were used for processing fluorescence signals, calculating growth rates, fitting exponential models, and generating all figures. The complete analysis code and datasets will be available in the public data repository on Zenodo upon publication.

